# The power and limits of predicting exon-exon interactions using protein 3D structures

**DOI:** 10.1101/2024.03.01.582917

**Authors:** Jeanine Liebold, Aylin Del Moral-Morales, Karen Manalastas-Cantos, Olga Tsoy, Stefan Kurtz, Jan Baumbach, Khalique Newaz

## Abstract

Alternative splicing (AS) effects on cellular functions can be captured by studying changes in the underlying protein-protein interactions (PPIs). Because AS results in the gain or loss of exons, existing methods for predicting AS-related PPI changes utilize known PPI interfacing exon-exon interactions (EEIs), which only cover ∼5% of known human PPIs. Hence, there is a need to extend the existing limited EEI knowledge to advance the functional understanding of AS. In this study, we explore whether existing computational PPI interface prediction (PPIIP) methods, originally designed to predict residue-residue interactions (RRIs), can be used to predict EEIs. We evaluate three recent state-of-the-art PPIIP methods for the RRI- as well as EEI-prediction tasks using known protein complex structures, covering ∼230,000 RRIs and ∼27,000 EEIs. Our results provide the first evidence that existing PPIIP methods can be extended for the EEI prediction task, showing F-score, precision, and recall performances of up to ∼38%, ∼63%, and ∼28%, respectively, with a false discovery rate of less than 5%. Our study provides insights into the power and limits of existing PPIIP methods to predict EEIs, thus guiding future developments of computational methods for the EEI prediction task. We provide streamlined computational pipelines integrating each of the three considered PPIIP methods for the EEI prediction task to be utilized by the scientific community.

## 1 Introduction

### 1.1 Background and motivation

Proteins physically interact with each other through protein-protein interactions (PPIs) to perform cellular functions. The collection of PPIs (i.e., the PPI interactome) changes with varying cellular conditions induced via several biological phenomena, including, e.g., alternative splicing (AS) [1]. AS produces multiple protein isoforms by joining the protein-coding regions (i.e., exons) of a gene in different combinations. Abnormal protein isoforms have been shown to adversely affect human health [2–4], and thus studying AS-related PPI changes is important. Note that, in this study, the term “PPI change” stands for the event that any two proteins forming a PPI in one cellular condition (e.g., healthy) do not form a PPI in another cellular condition (e.g., disease), or vice versa.

Using wet lab experiments to study PPI changes is expensive and time-consuming [5]. Hence, computational methods to study AS-related PPI changes were recently developed [6,7], which rely on existing knowledge about protein 3-dimensional (3D) structural domain-domain interactions (DDIs). One such recent method is ALTernatively spliced INteraction prediction (ALT-IN) [6], which extracts biochemical or DDI features of proteins to train a machine-learning model based on ground truth knowledge about existing AS-affected PPIs [1]. Then, ALT-IN uses the trained model to predict whether or not a PPI disappears when, for any of the participating proteins, the original isoform is replaced by an alternative isoform. While ALT-IN indirectly relies on DDIs, other methods use DDIs more directly to predict AS-affected PPIs. The general approach of such methods is to i) integrate existing knowledge about DDIs to first create a global DDI-resolved PPI network (i.e., a PPI network with information on which domains of the two proteins interact) and ii) use the global DDI-resolved PPI network to predict whether a PPI disappears if any of the two participating proteins lose the PPI-specific domain(s) due to AS [7]. One such recent method is Domain Interaction Graph Guided Explorer (DIGGER) which not only created a DDI-resolved PPI network but also delved deep into characterizing an exon-level view of PPIs [8], based on the following premise. Because many AS events remove or add exons to form protein isoforms, looking at exon-exon interactions (EEIs) in PPIs is more informative than just PPIs. Thus, EEIs can help more accurately explore AS-related PPI changes and their effect on cellular functions. This exon-level view of PPIs from DIGGER has been subsequently used by a recent method called NEASE to predict PPIs and functional changes due to AS in several biological conditions [9]. Additionally, the DIGGER data was incorporated into a recent AS-related PPI prediction method called LINDA to get a mechanistic understanding of AS-related PPI changes [10].

Thus, existing studies show that the knowledge about the exon-level view of PPIs capturing EEIs can help to predict AS-related PPI changes, which can then be used to understand the underlying molecular mechanisms. However, all of the existing AS-related PPI prediction methods rely on the existing EEI knowledge, which, although highly confident, covers only ∼5% of all known human PPIs [11]. Therefore, to fully capture the PPI network-based functional effects of AS, there is a need to extend the current limited knowledge of EEIs to the entire human proteome.

One way to extend the collection of EEIs is to predict novel EEIs using existing PPI interface prediction (PPIIP) methods [12], which can be grouped into two major categories. The first category of PPIIP methods takes a single protein as input and predicts all possible PPI interfaces (or binding sites) of the protein [13–16]. As is, this category of PPIIP methods cannot be trivially applied to the EEI prediction task. This is because even if one could use such methods to identify PPI interfacing exons of a protein, it is not trivial to find which exons of the protein’s PPI partners the identified interfacing exons interact with. The second category of PPIIP methods, which is more suitable for the EEI prediction task, takes two proteins of a PPI (but lacking the corresponding interface information) as input and predicts which parts of the two proteins interact, i.e., form a PPI interface [17]. Henceforth, whenever we use the term “PPIIP method”, we mean the PPIIP method from the second category. Typically, a PPIIP method uses ground truth knowledge about interacting (or non-interacting) residue pairs of PPI interfaces and protein sequence- or 3D structure-based features of residues to train a supervised machine-learning model. Such a trained model intuitively captures (dis)similarities between interacting or non-interacting residue pairs. Given any two test proteins *p_1_*and *p_2_* that are known to form a PPI but lack the corresponding interface knowledge, the trained model predicts the likelihood of any two residues *r_1_* from protein *p_1_* and *r_2_* from protein *p_2_* to interact, i.e., to form a residue-residue interaction (RRI) [17].

Motivated by the recent considerable increase in high-quality 3D protein structures [18,19], several deep learning-based PPIIP methods have been developed. Such methods take 3D protein structures as input, train a deep learning model using 3D protein structures or physicochemical-based features of residues, and then, given two proteins of a PPI with an unknown PPI interface, predict RRIs [20–25]. Three such recent PPIIP methods are Differentiable Molecular Surface Interaction Fingerprinting (dMaSIF) [20], Protein Interface Network (PInet) [21], and Graph learning of inter-protein contacts (GLINTER) [22]. While both dMaSIF and PInet use deep geometric learning [26], GLINTER relies on graph convolutional networks and residual neural networks [27]. While the three methods have not been compared to each other until now, at least one of them has been shown to outperform at least one of the six other existing PPIIP methods (Table 1), i.e., Dynamic Graph CNN [28], PointNet++ [29], BIPSPI [30], DeepHomo [24], PAIRpred [31], and ComplexContact [32]. Hence, dMaSIF, PInet, and GLINTER can be considered as state-of-the-art PPIIP methods.

**Table 1.** Summary of the existing PPIIP methods (columns) to which either dMasIF, PInet, or GLINTER (rows) were compared in their respective studies. Table cells with checkmarks indicate cases where dMaSIF, PInet, or GLINTER were compared against and outperformed the corresponding method in the marked column. Table cells with no checkmarks indicate that the corresponding pair of methods were not compared to each other.

|  | Dynamic Graph CNN [28] | PointNet++ [29] | BIPSPI [30] | PAIRpred [31] | Deep-Homo [24] | Complex-Contact [32] |
| --- | --- | --- | --- | --- | --- | --- |
| dMaSIF[20] | ✓ | ✓ |  |  |  |  |
| Plnet[21] |  |  | ✓ | ✓ |  |  |
| GLINTER[22] |  |  | ✓ |  | ✓ | ✓ |

Rather than using the PPIIP methods to predict EEIs, one could use docking [33], which aims to find an optimal spatial arrangement of any two 3D structures of proteins based on some predefined optimization criteria, e.g., minimizing the average distance of every residue pair between the two proteins. However, docking has high computational costs, taking a few minutes to several hours or even days to process one protein pair [33,34]. Given the estimated ∼650,000 possible human PPIs [35], using docking to predict EEIs over the entire estimated human PPIs would need at least ∼1.2 years if one protein pair is processed per minute. Similar to docking, the latest developments in protein complex predictions, such as AlphaFold-multimer [36], which take protein sequences as input and predict the 3D structure of the corresponding protein complex, typically take hours to process one protein pair [37] and hence do not scale well. In contrast to docking or the recent deep learning-based protein complex prediction methods, the existing PPIIP methods (mentioned above) have been specifically designed for the PPI interface predictions and are much more computationally efficient, most of which take only a few seconds to minutes to predict interfacing residues of a protein pair [20–22]. However, it has not been evaluated whether, or how well, the existing PPIIP methods can be used for the EEI prediction task, which is what we do in this work.

### 1.2 Our contributions

We evaluate three state-of-the-art PPIIP methods, i.e., dMaSIF, GLINTER, and PInet, in the EEI prediction task (Figure 1). Note that another PPIIP method called ProteinMAE [25] was recently published that learns protein surface representations with a masked autoencoder and self-supervised learning using all existing protein structural data, and outperformed dMaSIF. However, we could not use ProteinMAE in our current study because the corresponding source code is not publicly available (last checked on 29th Feb. 2024). We perform our study using all reviewed UniProt human proteins which map to ∼13,000 high-confidence co-crystallized 3D structures of protein pairs from the Protein Data Bank (PDB) [19], covering ∼230,000 interfacing RRIs and ∼27,000 interfacing EEIs (see Section 2.1 for details).

**Figure 1:**
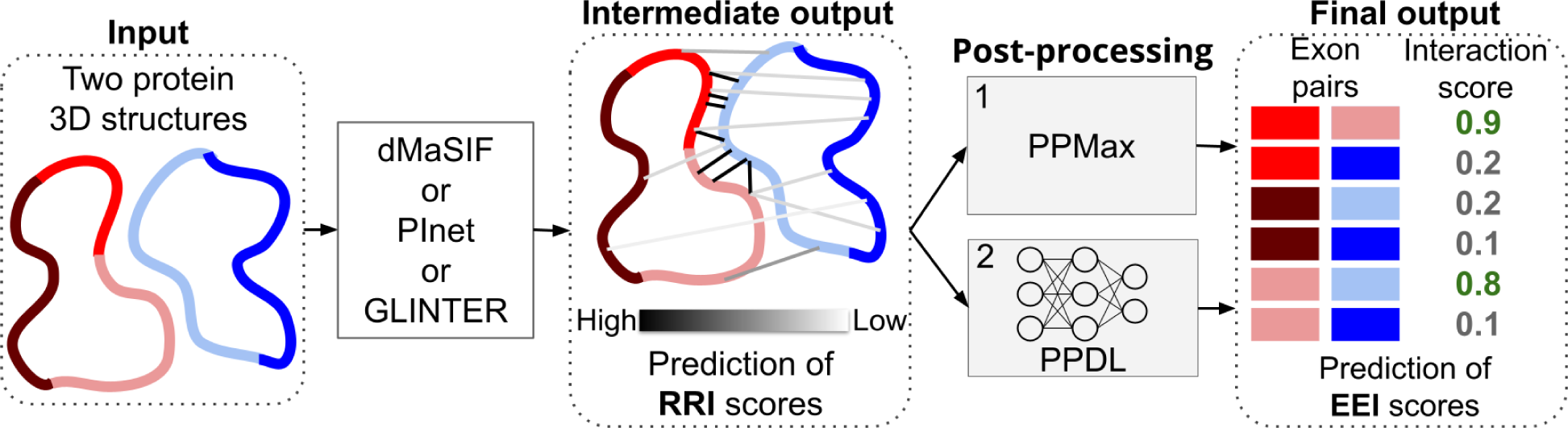
Our EEI prediction pipeline. Given the 3D structures of two proteins (represented in red and blue color in the extreme left of the figure) with exon annotations (highlighted in different intensities of red or blue) as input, we apply a PPIIP method (either dMaSIF, PInet, or GLINTER) to obtain RRI scores (indicated by lines of different intensities in the middle panel) for each pair of residues in the two proteins (”Intermediate output”). Given the RRI scores, we then obtain the final EEI scores (“Final output”) using two post-processing approaches, i.e., PPMax and PPDL (see Section 2.3 for details).

Given the considered PPIIP methods and data, we do the following. First, we evaluate the methods on the RRI prediction task for which the methods were originally designed. We do this because the three methods have never been compared to each other which allows us to establish a benchmark for their performance within their designated scope. Additionally, this would help us to better interpret their performance on the proposed EEI prediction task. For a robust performance evaluation, for each of the three methods, we perform 5-fold cross-validation [38] across three datasets (see Section 2.1 for details). Thus, in total, we train and test 3 × 5 × 3 = 45 machine learning models. Second, we evaluate the methods on the EEI prediction task for which we adapt them using two post-processing approaches. Similar to RRI prediction performance evaluations, for each of the three methods, for each of the three ways of defining ground truth EEIs, and for each of the two post-processing approaches to predict EEIs, we perform 5-fold cross-validation. Thus, in total, we train and test 3 × 3 × 2 × 5 = 90 machine learning models. We evaluate each method based on the area under the receiver operating characteristic (AU-ROC^1^) curve, the area under the precision-recall curve (AU-PRC), as well as precision, recall, and F-score for a certain false discovery rate (FDR).

Our key finding is that it is possible to extend the current 3D structure-based PPIIP methods to the task of EEI predictions. For both RRI and EEI predictions, each method performs significantly better than random with AU-ROC values reaching up to 85% for RRI predictions and 70% for EEI predictions. While the results vary with the choice of the dataset, significantly, for the EEI predictions, the maximum F-score, precision, and recall reach up to ∼38%, ∼63%, and ∼28%, respectively, with an FDR of less than 5%. Our study provides insights into how one can use existing 3D protein structure-based PPIIP methods to predict EEIs, and thus can guide future developments of computational methods for the EEI prediction task. Additionally, we provide a computational pipeline that streamlines the use of each of the three considered PPIIP methods for both the RRI and the EEI prediction tasks that can be utilized by the research community.

## 2 Data and methods

### 2.1 Dataset curation

We collect all 20,378 reviewed human proteins from UniProt [39] (downloaded Jan. 2022). We map the 20,378 UniProt protein sequences to the 3D structural data of co-crystallized human proteins from the PDB (downloaded on Jan. 2022), which results in 5,977 PDB entries with sufficient 3D structural resolution. We map exon information onto UniProt protein sequences using annotations from Ensembl [40], and we map 3D structural information from PDB onto UniProt protein sequences using SIFTS [41,42] (Section 2.1.1). Given a co-crystallized protein pair, we define the corresponding RRIs and EEIs (Section 2.1.2) using three PPI interface definitions to obtain three datasets (Figure 2). More details are outlined below and in Supplementary Section 1.1.

**Figure 2:**
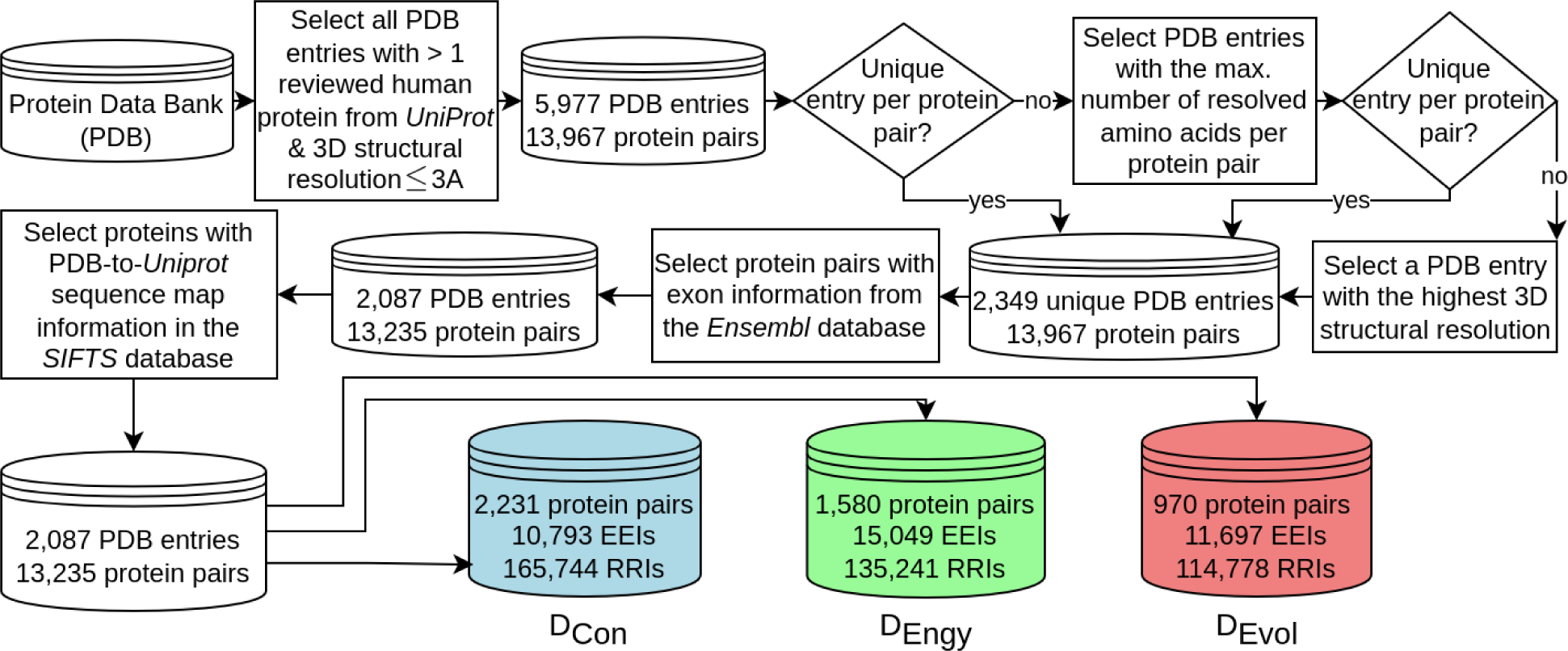
Overview of our data generation pipeline. We use all available confident PDB data to create three datasets, i.e., D_Con_, D_Engy_, and D_Evol_, for our study. See text for details.

#### 2.1.1 Protein co-crystal data

We consider a PDB entry if it has a 3D structural resolution of less than or equal to 3 Å and contains at least one co-crystallized human protein pair. This results in 2,349 unique PDB entries, which contain 13,967 unique co-crystallized human protein pairs involving 2,882 unique human proteins. For each of the 2,882 proteins, we download their amino acid sequences from the UniProt database. We only keep those proteins to which we could map the exon information according to the Ensembl database [40] using the file <u>Homo_sapiens.GRCh38.105.gtf.gz</u>. This results in 2,088 unique PDB entries that contain information about the co-crystallization of 13,240 (out of the 13,967) protein pairs, comprised of 2,660 unique proteins. From these proteins, we select those for which we could map their sequence positions onto their 3D resolved structures from the corresponding PDB entries using the SIFTS database. This results in 2,649 proteins from 2,064 PDB entries with information about the co-crystallization of 13,190 unique protein pairs. This is the final set of unique proteins, PDB entries, and co-crystallized protein pairs that we use in our study.

#### 2.1.2 Definition of RRIs and EEIs

Given a pair of co-crystallized proteins *p_1_* and *p_2_*, we define an RRI between residues *r_1_* from *p_1_*and *r_2_* from *p_2_*, if and only if the 3D Euclidean distance between any of the heavy atoms (i.e., Carbon, Nitrogen, Oxygen, and Sulphur) of *r_1_* is within 6 Å to any of the heavy atoms of *r_2_*. This definition of RRIs is consistent with existing studies [22,30,43–46].

For each pair of exons *e_1_* from *p_1_* and *e_2_* from *p_2_*, we evaluate whether the two exons form an EEI using three established PPI interface detection methods, as outlined below.

##### Contact-based (D_Con_)

We define an EEI between two exons *e_1_* and *e_2_* if and only if at least one residue of *e_1_* interacts with at least one residue of *e_2_*. We define an interaction between a pair of residues in the same manner as we do in the previous paragraph. Among 2,064 PDB entries, we only consider those 1,543 entries that i) contain at least one co-crystallized protein pair with at least one interacting exon pair and ii) could be preprocessed by each of the three PPIIP methods (Section 2.2). Among 13,190 co-crystallized protein pairs, 2,231 pairs remain that contain 10,793 interacting and 38,060 non-interacting exon pairs. Among the 10,793 EEIs, we obtain 165,744 interacting and 62,216,726 non-interacting residue pairs.

##### Energy-based (D_Engy_)

We use Protein Interfaces, Surfaces, and Assemblies (PISA) [47] to identify biologically relevant interfaces of co-crystallized proteins. PISA first computes a solvation free energy gain of an interface, which quantifies the thermodynamic changes that occur due to the interface formation. Then it computes the probability of observing the same solvation free energy gain by chance when a random set of atoms (with the same area as that of the interface) is picked from the non-interfacing surfaces of the two proteins. PISA labels an interface as biologically relevant if the corresponding probability is less than 0.5. We define an EEI between *e_1_* and *e_2_* if and only if they overlap with biologically relevant interfaces. Among 2,064 PDB entries, we only consider those 1,321 entries that i) contain at least one co-crystallized protein pair with at least one interacting exon pair and ii) could be preprocessed by each of the three PPIIP methods (Section 2.2). Among 13,190 co-crystallized protein pairs, 1,580 protein pairs remain that contain 15,049 interacting and 28,227 non-interacting exon pairs. Among the 15,049 EEIs, we obtain 135,241 interacting and 54,918,541 non-interacting residue pairs.

##### Evolution-based (D_Evol_)

We use the Evolutionary Protein-Protein Interface Classifier (EPPIC) [48] to identify biologically relevant interfaces of co-crystallized proteins. EPPIC characterizes an interface as biologically relevant if the interface surface area is more than 2,200 Å^2^, while it characterizes an interface to be an artifact of crystallization if the interface surface area is less than 400 Å^2^. For interface surface areas between 400 Å^2^ and 2,200 Å^2^, EPPIC uses information about the evolutionary conservation of the corresponding protein sequences to characterize whether the interface is biologically relevant or not. For our study, we run EPPIC using default parameters. We define an EEI between *e_1_* and *e_2_* to interact if and only if they overlap with biologically relevant interfaces. Among 2,064 PDB entries, we only consider those 898 entries that i) contain at least one co-crystallized protein pair with at least one interacting exon pair and ii) could be preprocessed by each of the three PPIIP methods (Section 2.2). Among 13,190 co-crystallized protein pairs, 970 protein pairs remain that contain 11,697 interacting and 20,710 non-interacting exon pairs. Among the 11,697 EEIs, we obtain 114,778 interacting and 42,317,056 non-interacting residue pairs.

### 2.2 PPI interface prediction methods

We use three recently established PPIIP methods, i.e., dMaSIF, GLINTER, and PInet, for reasons outlined in Section 1. Given two protein 3D structures that are known to form a PPI, a PPIIP method either predicts interactions between pairs of atoms, residues, or structural regions of the two proteins. We use each of the three methods to predict RRI scores for each pair (*r_1_*, *r_2_*) of residues between two input proteins *p_1_* and *p_2_*, where *r_1_* is from *p_1_* and *r_2_* is from *p_2_*. A higher RRI score indicates a higher likelihood of interaction between the corresponding residue pair (see Supplementary Section 1.2 for details). Note that, unless stated otherwise, we run all methods with their recommended parameter values to preprocess the input files and train the corresponding model.

### 2.3 Post-processing approaches for EEI prediction

Given two proteins *p_1_* and *p_2_* and the corresponding RRI predictions from a PPIIP method, we propose two post-processing approaches to predict the EEI scores between any two exons *e_1_*from *p_1_* and *e_2_* from *p_2_*. The first post-processing approach (referred to as post-processing via the maximum RRI score or *PPMax* for short) calculates the score for an exon pair (*e_1_,e_2_*) as the maximum predicted RRI score among all residue pairs (*r_1_*,*r_2_*), where *r_1_* is a residue in exon *e_1_* and *r_2_* is a residue in exon *e_2_*. The second post-processing approach (referred to as post-processing via deep learning or *PPDL* for short) is based on a convolutional neural network [49] where, given a set of interacting and non-interacting exon pairs, it first trains a model to learn distinguishing patterns of the predicted RRI scores for interacting vs. non-interacting exons. Then, given two test exon pairs, it predicts the corresponding EEI score. More details on PPDL are described next.

To create inputs for PPDL, given a PPIIP method, for each pair of exons *e_1_* and *e_2_* with *m* and *n* residues respectively, we use the PPIIP method to obtain an *m* × *n* matrix *M*(*e_1_*,*e_2_*). An entry *M*(*e_1_*,*e_2_*)[*i*,*j*] is the predicted RRI score of the *i*^th^ residue in *e_1_* and the *j*^th^ residue in *e_2_*. We focus on only those pairs (*e_1_*,*e_2_*) such that the number of residues in *e_1_* and the number of residues in *e_2_* are not larger than 100. For each of our datasets (Section 2.1.2), this selection criterion is satisfied by more than 95% of the exon pairs. We keep 10,091 interacting and 37,233 non-interacting exon pairs for D_Con_, 14,530 interacting and 27,889 non-interacting exon pairs for D_Engy_, and 11,301 interacting and 20,454 non-interacting exon pairs for D_Evol_ (Supplementary Table S1). The input to our PPDL approach is a *100*×*100*-matrix *M*(*e_1_*,*e_2_*) corresponding to each exon pair (*e_1_*,*e_2_*). For exons shorter than 100, we use zero padding. The output of PPDL is a continuous score ranging from 0 to 1 where a higher score means a higher likelihood of interaction. Our PPDL architecture consists of two convolutional layers, two max-pooling, two non-linear activations for feature extraction, and two linear layers for the final classification (Supplementary Figure S1).

### 2.4 Model training and evaluation

For each dataset (Section 2.1.2), we perform RRI or EEI predictions as follows. We split the set of protein pairs in the dataset randomly into five disjoint subsets. Four of the subsets contain *n*/5 (rounded up to the nearest integer) protein pairs, while the fifth subset contains the remaining protein pairs. We use these subsets in a 5-fold cross-validation manner. That is, we use each subset as test data and the remaining four subsets as the training data. We use the same five subsets for RRI and EEI predictions across all three PPIIP methods.

For RRI predictions, the protein pairs in the training data are used to train a model, which is then used to predict RRI scores between protein pairs in the corresponding test data. This way, we train five models of RRI predictions for a given method and dataset combination. Because we use three different datasets and PPIIP methods, with 5-fold cross-validation, we train and test 3×3×5=45 RRI prediction models.

For EEI predictions, given a PPIIP method, we first use a model trained for RRI predictions to obtain the RRI scores for each protein pair in the corresponding test data. We do this for each of the five trained models, which gives RRI scores for each protein pair in the entire dataset. Then we use the predicted RRI scores to obtain the EEI scores using two post-processing approaches, as outlined in Section 2.3. Specifically, for PPMax-based EEI predictions, we take the maximum RRI score between a pair of exons to obtain the corresponding EEI score, without training an additional machine learning model. For PPDL-based EEI predictions, we use the RRI scores of the exon pairs in the training data to train an additional model (on top of the PPIIP method from which we obtain the corresponding RRI scores) and then test the trained model using the RRI scores of exon pairs in the test data. Because we use three different datasets, three PPIIP methods, and two post-processing approaches, with 5-fold cross-validation, we train and test 3×3×2× 5=90 EEI prediction models.

We evaluate each of the trained models using five performance measures: AU-ROC, AU-PRC, precision, recall, and F-score. While AU-ROC and AU-PRC can be computed over all prediction scores of the test data, the computation of precision, recall, and F-score requires a predefined score threshold (i.e., “decision threshold”), such that residue (or exon) pairs with scores greater than the decision threshold are predicted to be interacting and the residue (or exon) pairs with scores less than or equal to the decision threshold are predicted to be non-interacting. We define such a decision threshold as follows. Given a test set, we take the predictions of all non-interacting residue (or exon) pairs from the training set as our background (or null) distribution of non-interacting residue (or exon) pair scores (Supplementary Figure S2). Then, we define a decision threshold as the score that accepts *t*% of non-interacting residue (or exon) pairs (i.e., false positives) from this background with *t* varying from 1 to 5 in increments of 1, which corresponds to the FDRs of 1-5% in increments of 1%.

## 3 Results and discussion

We evaluate three state-of-the-art 3D structure-based PPIIP methods, i.e., dMaSIF, PInet, and GLINTER, on how well the methods can predict EEIs between two proteins. The considered PPIIP methods were originally designed to predict RRIs and not EEIs, but were, to the best of our knowledge, never compared to each other. Hence, first, we compare them on the RRI prediction task to see whether any of them performs better than the other two (Section 3.1). Second, to adapt the PPIIP methods for the EEI prediction task, we propose two post-processing approaches, which we compare against each other to select the best one for further analyses (Section 3.2). Third, we compare the considered PPIIP methods for the EEI prediction task using the best post-processing approach to evaluate how well we can predict EEIs using existing PPIIP methods (Section 3.3). Fourth, we compare the performance of the PPIIP methods on the RRI prediction task versus the EEI prediction task to evaluate how well the methods predict interactions between proteins at the two different abstraction levels (i.e., residues vs. exons, Section 3.4). Finally, we evaluate the computational runtime of the PPIIP methods to relate their performance accuracies with their computational efficiencies (Section 3.5).

### 3.1 dMaSIF performs the best in the RRI prediction task

For each dataset, namely D_Evol_, D_Engy_, and D_Con_ (Section 2.1.2), each method performs non-randomly with average (over the five cross-validation folds) AU-ROC values ranging from 0.62 to 0.87 (Figure 3a and Supplementary Table S2). dMaSIF consistently performs best, with average AU-ROC values of 0.87 for D_Evol_, 0.86 for D_Engy_, and 0.83 for D_Con_, suggesting that the AU-ROC-based performance of dMaSIF is robust across datasets. The performance of GLINTER closely follows dMaSIF, with average AU-ROC scores of 0.84 for D_Evol_, 0.83 for D_Engy_, and 0.76 for D_Con_. PInet performs far worse than both dMaSIF and GLINTER across all datasets, with average AU-ROC scores of 0.68 for D_Evol_, 0.65 for D_Engy_, and 0.62 for D_Con_.

**Figure 3:**
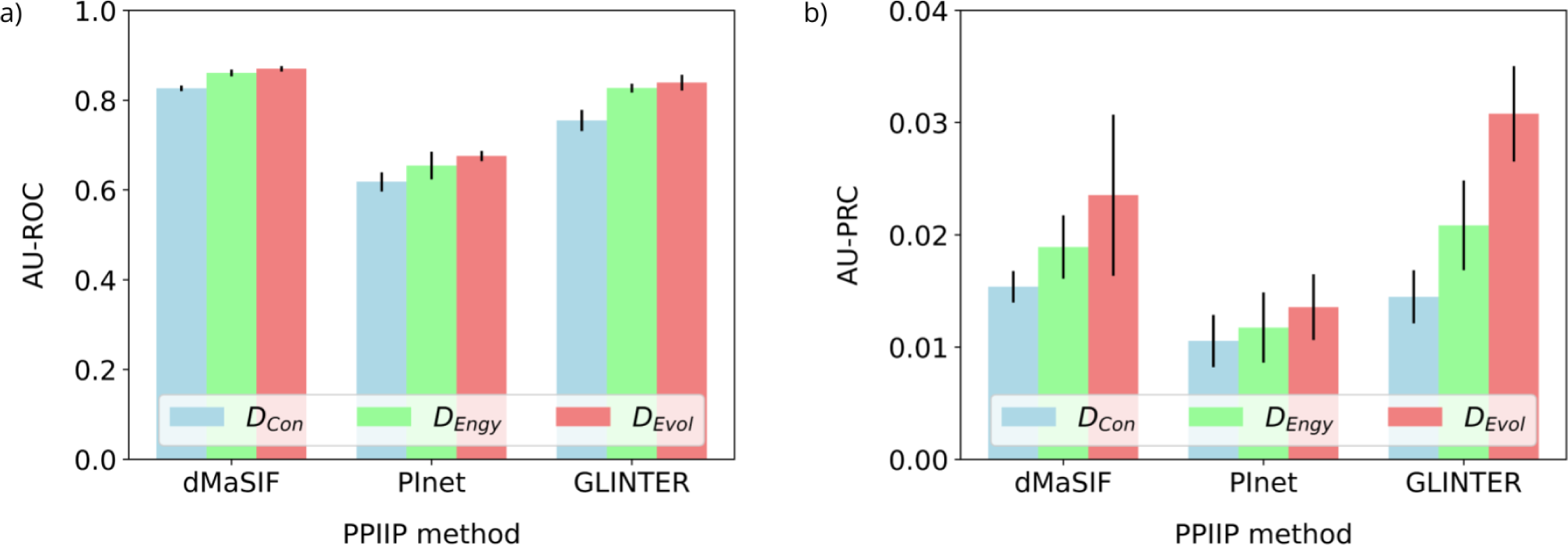
Performances of the PPIIP methods on the RRI prediction task based on AU-ROC (panel a) and AU-PRC (panel b) for each of the three datasets (indicated by the three colors). In a panel, the height of a given bar represents the mean AU-ROC (or AU-PRC) score across the five cross-validation folds and the vertical line above the bar indicates the corresponding standard deviation.

Although AU-ROC is commonly used to evaluate the performances of PPIIP methods in the RRI prediction task, it is less useful in our application, due to the imbalance in RRI datasets which contain ∼350 times more negative RRIs than positive RRIs (Section 2.1.2). For such imbalanced data sets, AU-PRC gives a more accurate quantitative value of how well the methods predict interacting RRIs compared to each other [50]. We find that the average AU-PRC values are below 0.04 for each of the PPIIP methods across each of the three datasets (Figure 3b). Note that because we use state-of-the-art PPIIP methods, the low AU-PRC values indicate that the underlying problem of RRI predictions is far from being solved and novel improved PPIIP methods are required. Overall, for D_Con_, independent of whether we use AU-ROC or AU-PRC, the ranking of methods remains the same, i.e., dMaSIF performs best, followed by GLINTER and PInet. However, based on AU-PRC, for datasets D_Engy_, and D_Evol_, GLINTER performs best, followed by dMaSIF and PInet, a ranking which differs from the one based on AU-ROC.

In addition to AU-ROC and AU-PRC, we measure the methods’ performances using precision, recall, and F-score at a prediction score decision threshold that allows for an FDR of *t*% (see Section 2.4 for details). To test the robustness of results across different decision thresholds, we vary the value of *t* from 1 to 5 in increments of 1. Given an FDR, we perform 45 performance tests for each PPIIP method corresponding to the combinations of the three performance measures (i.e., precision, recall, and F-score), the three datasets, and the five cross-validation folds.

We find that dMaSIF performs best for the majority of the performance tests for each FDR choice except 1%, followed by GLINTER and PInet (Figure 4a). For the 1% FDR, GLINTER performs almost equally well as dMaSIF, followed by PInet. Additionally, for each of precision, recall, and F-score, we evaluate whether one method significantly outperforms the other two using the paired Wilcoxon rank test. That is, given a pair of methods and a performance measure for a given FDR, we compare their performance scores across all combinations of the three datasets and the five cross-validation folds (15 tests in total). For each performance measure, dMaSIF significantly (q-value < 0.05) outperforms at least one of PInet or GLINTER for most of the FDR choices, GLINTER significantly (q-value < 0.05) outperforms PInet for the FDR choice of 5%, while PInet never outperforms any of the other two PPIIP methods (Supplementary Table S3).

**Figure 4:**
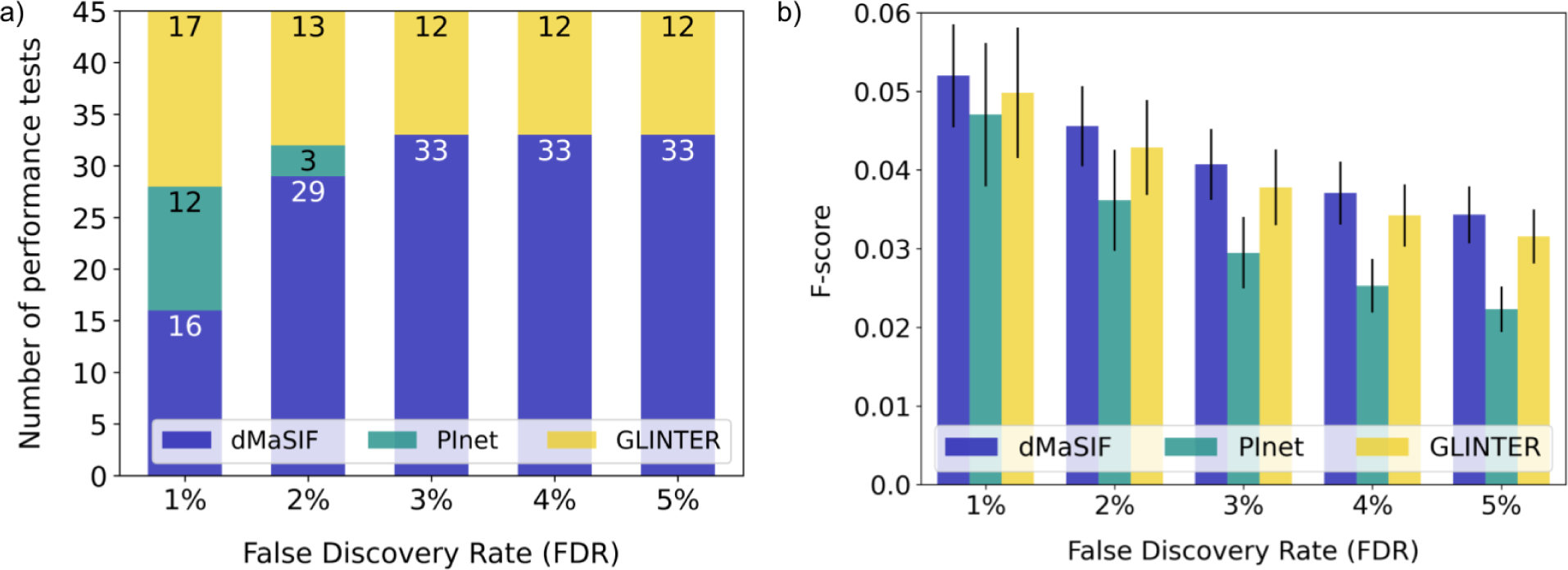
Performances of the PPIIP methods on the RRI prediction task. Panel a shows the distribution of the number of performance tests in which a given method performs best. The x-axis shows the FDR and the y-axis the number (45 in total) of performance tests. For an FDR value, a bar shows the number of performance tests in which the methods perform best (indicated by different colored compartments of the bar and the corresponding numbers within those compartments). For example, for the 1% FDR, dMaSIF, PInet, and GLINTER perform the best in 16, 12, and 17 performance tests, respectively. Panel b shows the F-score of the PPIIP methods for the D_Engy_ dataset. Results are qualitatively similar for the D_Evol_ dataset. For the D_Con_ dataset, although the overall trend is the same (i.e., there is a decrease in F-score with the increase in FDR), GLINTER performs the worst among the three PPIIP methods at each FDR choice (Supplementary Table S2). The height of a given bar represents the mean F-score across the five cross-validation folds and the vertical line above the bar indicates the corresponding standard deviation.

Looking at absolute F-score values at different FDR choices, we find that the performance values for each method across the different FDR choices are very low (Figure 4b). Note that there is a decrease (although marginal) in the F-score values with the increasing FDR. This is because as the FDR threshold increases, although the recall values of the PPIIP method increase, their precision values decrease drastically (Supplementary Table S2).

Given a PPIIP method, its performance varies across datasets. For AU-ROC and AU-PRC, any given method always performs best for D_Evol_ followed by D_Engy_, and D_Con_. For each of F-score (Figure 4b), precision, and recall averaged over the five decision thresholds and the five cross-validation folds, dMaSIF, PInet, and GLINTER perform the best for D_Evol_ followed by D_Engy_, and D_Con_, with one exception where PInet’s recall is the best for D_Engy_ followed by D_Evol_ and D_Con_ (Supplement Table S2). Overall, GLINTER shows the largest variation in performance across datasets. For example, for average AU-PRC over the five cross-validation folds, GLINTER’s performance for D_Evol_ is about twice as good as its performance for D_Con_. However, for dMaSIF and PInet, the corresponding performances are ∼1.5 and ∼1.3 times higher, respectively. This signifies that GLINTER’s performance is more sensitive to the choice of the dataset, while PInet and dMaSIF show more robust performances across datasets.

To summarize, while PInet shows lower performance variations across datasets, it always performs the worst irrespective of the choice of dataset or performance measure. For some combinations of datasets and performance measures, GLINTER shows better performance than dMaSIF, but in the majority of cases, dMaSIF performs best. The consistently lower performance of PInet than dMaSIF or GLINTER could be due to the underlying methodology of the individual PPIIP methods. That is, given two proteins, both dMaSIF and GLINTER were explicitly designed to identify pairs of residues across the two proteins that interact. In contrast, PInet was designed to identify pairs of regions across the two proteins that interact, without explicitly identifying the interacting residue pairs within those pairs of regions.

### 3.2 PPDL outperforms PPMax in the EEI prediction task

We compare the two post-processing methods (PPMax and PPDL) on the EEI prediction task based on each of the performance measures that we used in the previous section. Given a PPIIP method, for a majority of the combinations of performance measures and datasets, we observe clear advantages of using PPDL over PPMax (Figure 5).

**Figure 5:**
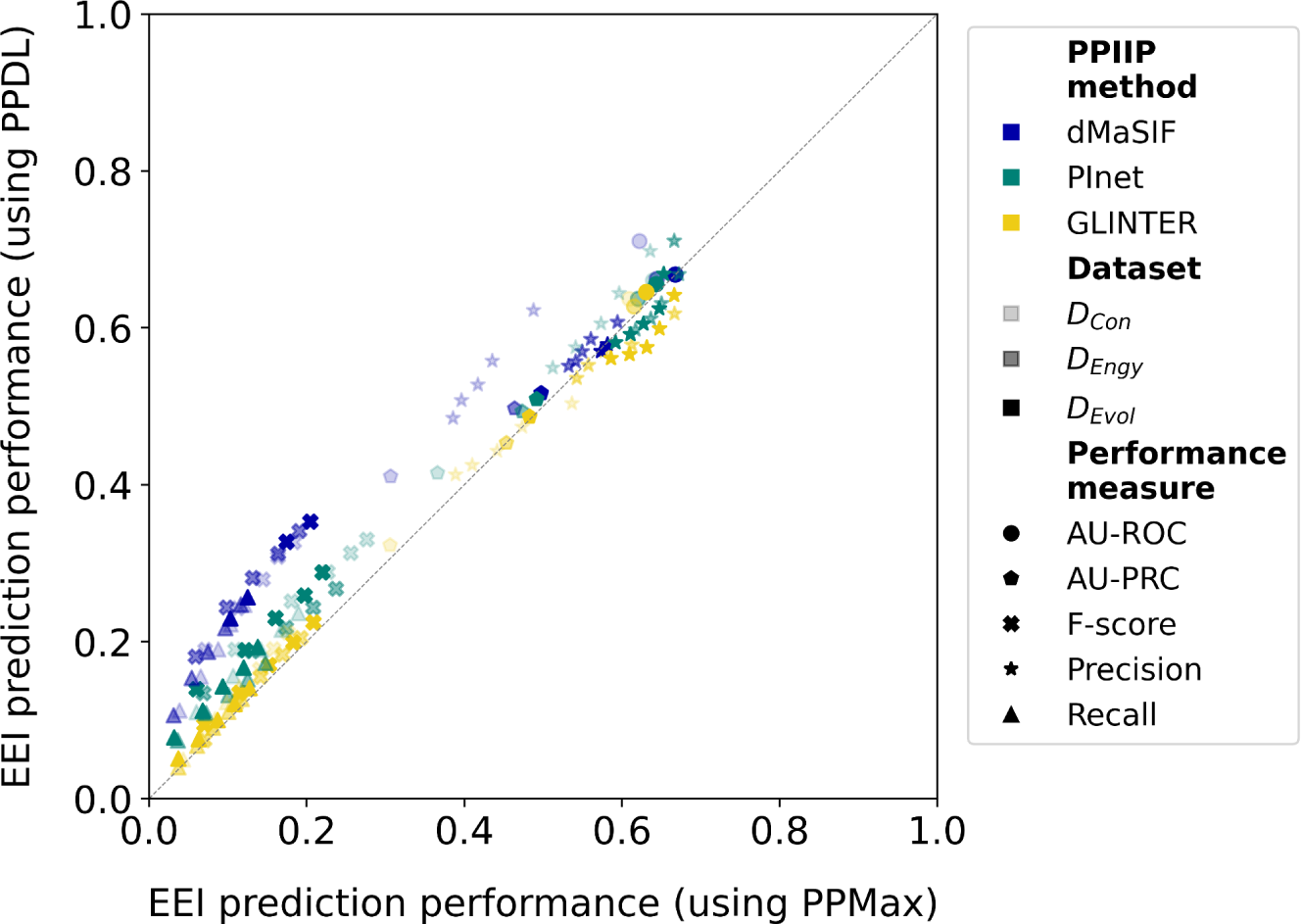
Performance of the PPIIP methods on the EEI prediction task using the PPMax (x-axis) versus the PPDL (y-axis) post-processing approaches. Each point in the plot represents the average (over the five cross-validation folds) performance of a PPIIP method using PPDL vs. using PPMax for a given combination of dataset, performance measure, and FDR choice (if applicable). For detailed results, see Supplementary Tables S4 and S5.

With respect to each of AU-ROC and AU-PRC, for each of the PPIIP methods, when compared across all 15 combinations of the three datasets and the five cross-validation folds, PPDL significantly (q-value < 0.05) outperforms PPMax (Supplementary Tables S3). With respect to precision, recall, and F-score, for a given PPIIP method, when compared across all 15 combinations of the three datasets and the five cross-validation folds at a given FDR, we find the following. For precision, for each of the PPIIP methods, although PPDL on average performs better than PPMax in most cases, the differences in the performances are non-significant (Supplementary Table S3). For recall and F-score, the performance significance of PPDL over PPMax varies depending on the PPIIP method and FDR choice. For dMaSIF, based on both recall and F-score, PPDL significantly (q-value < 0.05) outperforms PPMax at 4% and 5% FDR, while for the rest of the three FDR choices, the performance differences between PPDL and PPMax are non-significant (Supplementary Table S3). For each of PInet and GLINTER, based on both recall and F-score, PPDL significantly (q-value < 0.05) outperforms PPMax at each of the FDR choices (Supplementary Table S3).

These results signify the advantage of learning patterns (as in PPDL), rather than simply taking the maximum (as in PPMax), of the predicted RRI scores of two exons to predict whether the exons interact or not. Hence, in this manuscript, we only show detailed results for EEI predictions using PPDL, while we provide the corresponding results for PPMax in the supplement (Supplementary Table S4).

### 3.3 Existing PPIIP methods designed for RRI predictions can be extended to predict EEIs

For each dataset, namely D_Con_, D_Engy_, and D_Evol_ (Section 2.1.2), each method performs non-randomly with AU-ROC average (over the five cross-validation folds) values ranging from 0.63 to 0.71 (Figure 6a and Supplementary Table S5). dMaSIF consistently performs best, with average (over the five cross-validation folds) AU-ROC values of 0.71 for D_Con_, 0.66 for D_Evol_, and 0.66 for D_Engy_. PInet closely follows dMaSIF with average AU-ROC scores of 0.67 for D_Con_, 0.65 for D_Evol_, and 0.64 for D_Engy_. GLINTER performs the worst with average AU-ROC scores of 0.64 for D_Evol_, 0.63 for D_Con_, and 0.63 for D_Engy_. We find that the ranking of methods for the D_Engy_ and D_Evol_ datasets remains the same when we rank by AU-PRC instead of AU-ROC, i.e., dMaSIF performs best, followed by PInet and GLINTER (Figure 6b). However, unlike AU-ROC, considering AU-PRC for the D_Con_ dataset, PInet performs best, followed by dMaSIF and GLINTER.

**Figure 6:**
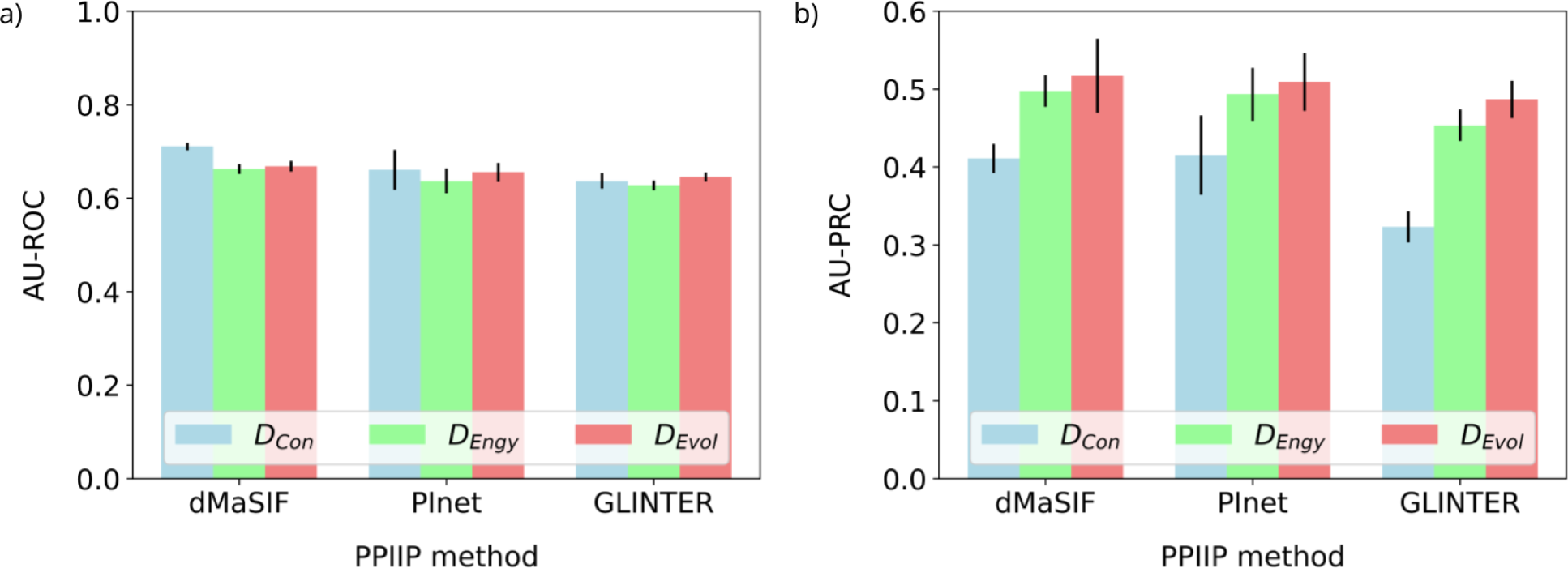
Performances of the PPIIP methods on the EEI prediction task based on AU-ROC (panel a) and AU-PRC (panel b) measures for each of the three datasets (indicated by the three colors). In a panel, the height of a given bar represents the mean AU-ROC (or AU-PRC) score across five-fold cross-validation and the vertical line above the bar indicates the corresponding standard deviation.

In addition to AU-ROC and AU-PRC, similar to how we evaluate the performance of the PIIP methods in the RRI prediction task, we measure the performances of the PPIIP methods in the EEI prediction task using precision, recall, and F-score at a decision threshold that allows for an FDR of *t*% (see Section 2.4 for details). Given an FDR, we perform 45 performance tests for each PPIIP method corresponding to the combinations of the three performance measures, the three datasets, and the five cross-validation folds.

We find that dMaSIF performs best for more than half of the tests for each FDR choice, followed by PInet and GLINTER (Figure 7a). To quantify the dominance of one method over the other two at a given FDR, for each of precision, recall, and F-score, we evaluate whether a method significantly outperforms the other two across all three datasets and five cross-validation folds using the paired Wilcoxon rank test. This approach is similar to how we compare the RRI prediction performance of the methods. For each performance measure, PInet significantly (q-value < 0.05) outperforms GLINTER for all FDR choices. PInet shows non-significant differences with dMaSIF with the following two exceptions. At the 4% FDR, PInet significantly outperforms dMaSIF based on precision. At the 4% and 5% FDR choices, dMaSIF significantly (q-value < 0.05) outperforms both PInet and GLINTER based on recall and F-score (Supplementary Table S3). For the rest of FDR choices of 1%, 2%, and 3%, dMaSIF does not show significant differences with either PInet or GLINTER based on any performance measure.

**Figure 7:**
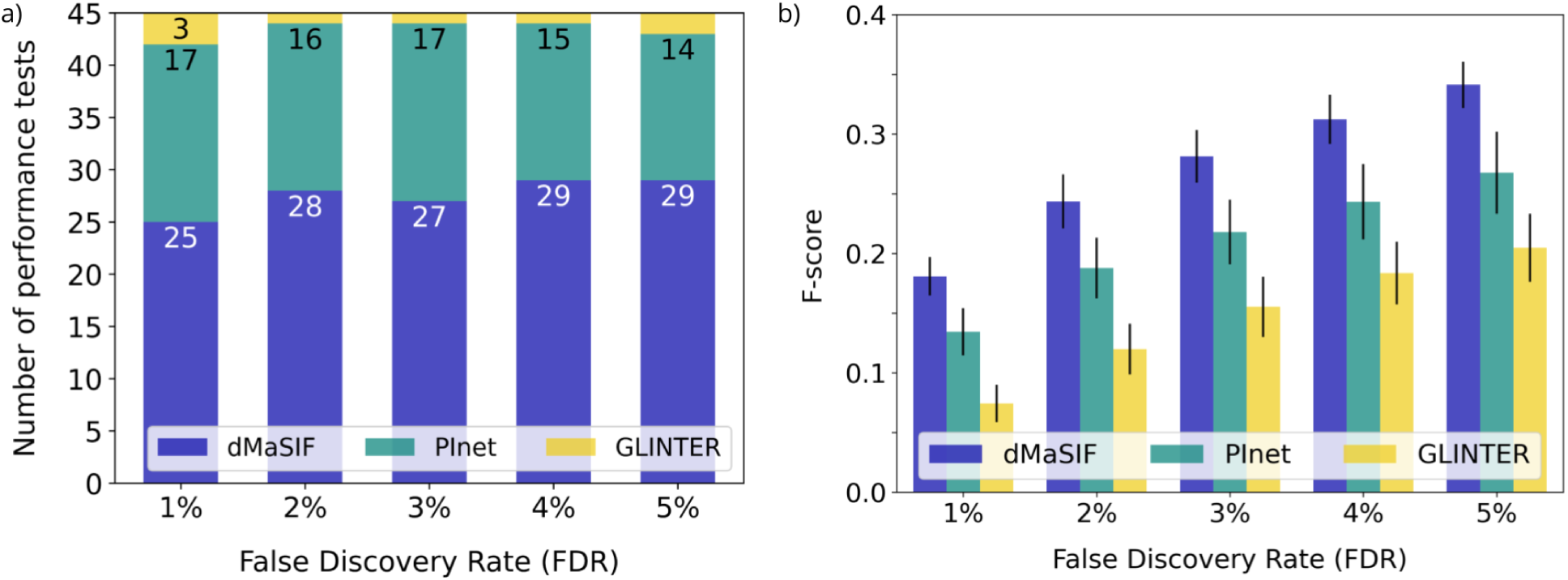
Performances of the PPIIP methods on the EEI prediction task. Panel a shows the distribution of the number of performance tests in which a method performs best. The figure can be interpreted in the same manner as Figure 4. Panel b shows the F-score of the PPIIP methods for the D_Engy_ dataset. Results are qualitatively similar for the D_Evol_ dataset. For the D_Con_ dataset, although the overall trend is the same (i.e., there is an increase in F-score with the increase in FDR), PInet performs equally as well as dMaSIF at each FDR choice (Supplementary Table S2). The height of a given bar represents the mean F-score across the five cross-validation folds and the vertical line above the bar indicates the corresponding standard deviation.

Looking at absolute F-score values at different FDR choices, we find that the performance values for each method increase with the increase in FDR (Figure 7b). This happens because as the FDR increases, for each method, we see a larger increase in recall with a much smaller drop in precision (Supplementary Table S2).

Similar to RRI predictions (Section 3.1), the EEI prediction performance of a PPIIP method varies across datasets. For AU-ROC, all methods perform equally well for all datasets, except for dMaSIF, which performs best for D_Con_. For AU-PRC, all methods always perform the best for D_Evol_ followed by D_Engy_ and D_Con_. For each F-score (Figure 7b), precision, and recall averaged over the five FDR choices and five cross-validation folds, dMaSIF and GLINTER perform the best for D_Evol_ followed by D_Engy_ and D_Con_, with some exceptions. That is, GLINTER’s recall and F-score as well as dMaSIF’s recall are better for the D_Con_ dataset than for the D_Evol_ dataset (Supplementary Table S5). Similarly, PInet performs best for D_Con_ with respect to F-score and recall, while it performs best for D_Engy_ with respect to precision. Although all methods show some variations in their performances across datasets, GLINTER shows the largest variation. For example, for average AU-PRC over the five cross-validation folds, GLINTER’s performance for D_Evol_ is 1.5 times larger than its performance for D_Con_, while for dMaSIF and PInet, their performance for D_Evol_ is 1.25 times larger than its performance for D_Con_.

To summarize, GLINTER not only shows the largest variation of its performance across the datasets, it also always performs the worst irrespective of the choice of dataset or performance measure. For some combinations of datasets and performance measures, PInet shows better performance than dMaSIF, but in the majority of cases, dMaSIF performs best. Hence, dMaSIF should be the preferred method of choice for the EEI prediction task.

### 3.4 Comparison of the PPIIP methods on the RRI vs. the EEI prediction tasks

To evaluate how the PPI interface prediction performances of the considered PPIIP methods vary across the two different abstraction levels, i.e., exons vs. residues, we compare the methods’ performances on the RRI vs. the EEI prediction tasks (Figure 8).

**Figure 8:**
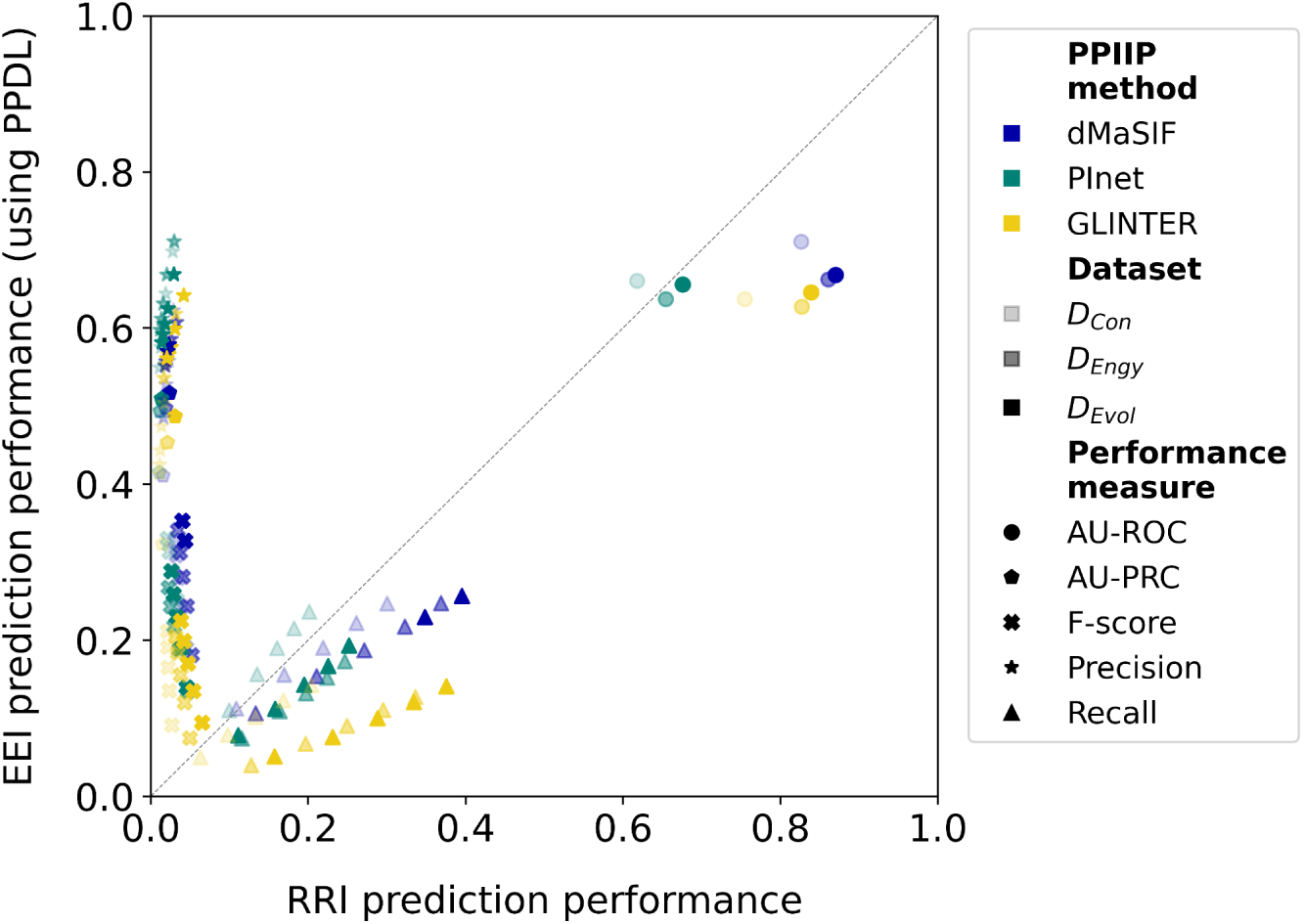
Performances of the PPIIP methods on the RRI (x-axis) vs. the EEI (y-axis) prediction tasks. A point in the plot represents the average (over the five cross-validation folds) performance of a method (indicated by color) on a given dataset (indicated by intensity) and a performance measure (indicated by the shape). For F-score, precision, and recall, results for all five FDR choices are shown. Hence, for a given method, there are 15 points for each of precision, recall, and F-score, which corresponds to the combination of three datasets and five FDR choices. For AU-ROC and AU-PRC, there is no FDR choice, and hence, given a method, for each of AU-ROC and AU-PRC, there are only three points corresponding to the three datasets.

For each of the PPIIP methods, EEI prediction performance is significantly (q-value < 0.05) better than RRI prediction performance based on AU-PRC, precision, and F-score, while RRI prediction performance is significantly (q-value < 0.05) better than EEI prediction performance based on recall (Supplementary Table S3). The differences in the precision of EEI vs. RRI predictions could be related to the differences in their respective ratios of positive and negative interactions. That is, the RRI datasets are highly imbalanced with ∼350 times more negative samples than positive samples, while the EEI datasets show much less data imbalance with only around twice as many negative samples than positive samples. This huge imbalance in the RRI datasets can make it challenging for the prediction models to correctly identify true interactions amidst a large number of negative instances. AU-ROC values for RRI prediction are better than those of EEI prediction for most PPIIP methods, except PInet where AU-ROC values for RRI and EEI predictions are comparable. Because AU-ROC quantifies how well a model distinguishes between true and false positives going from highest-ranking predictions to the lowest, it is less sensitive to class imbalance and hence gives a clearer picture of whether a method works better for the RRI vs. the EEI prediction task. Hence, for each PPIIP method, its RRI prediction performance is better than its EEI prediction performance.

### 3.5 Comparison of runtime

To evaluate the applicability of the trained EEI prediction models of the PPIIP methods for real-world usage on unseen data, we measure their computational runtime on a chosen set of 10 protein pairs (Supplementary Table S6). Given the protein pairs, we first measure the preprocessing runtimes of the PPIIP methods. dMaSIF, PInet, and GLINTER take on average (over the 10 protein pairs) ∼1.7 seconds (s), ∼36 s, and ∼359 s, respectively, to preprocess one protein pair. A much higher preprocessing runtime for GLINTER comes from the multiple sequence alignment step, which is not required for dMaSIF and PInet. Next, given the preprocessed protein pairs as input, we measure the prediction runtimes of the PPIIP methods. dMaSIF, PInet, and GLINTER take on average (over the 10 protein pairs) ∼2.9 s, ∼1.3 s, and ∼1.2 s, respectively, to predict the outcome for one protein pair (Table 2). Because dMaSIF has almost no preprocessing runtime, its total (preprocessing + prediction) runtime of ∼4.6 seconds is ∼78 and ∼8 times lower than the total time for GLINTER and PInet, respectively.

**Table 2.**
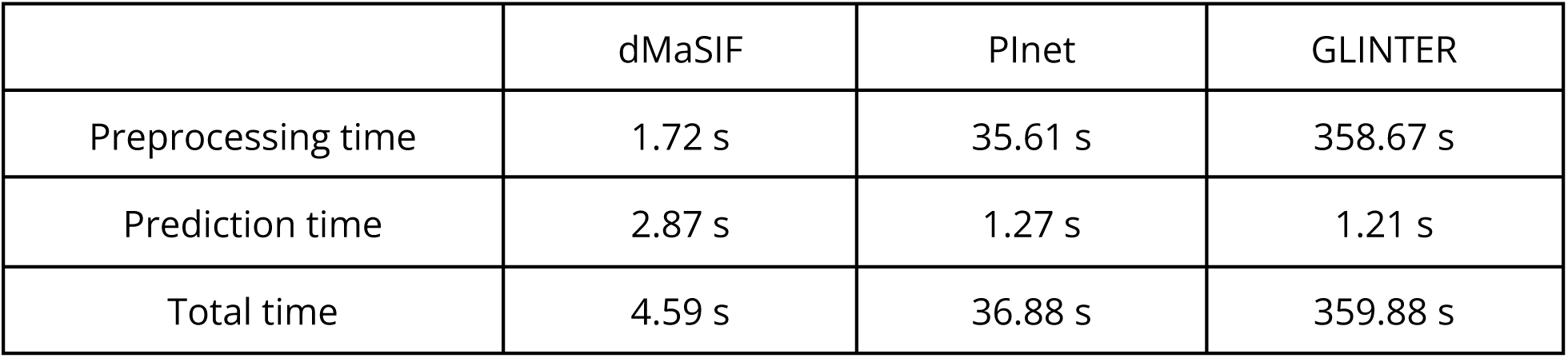
Average computational runtimes of the PPIIP methods per protein pair.

## 4 Conclusion

In this work, we examine whether existing PPIIP methods can be adapted to predict EEIs. We extend three state-of-the-art protein 3D structure-based PPIIP methods, i.e., dMaSIF, GLINTER, and PInet, via two post-processing approaches (PPMax vs. PPDL) that utilize RRI prediction scores of the PPIIP methods, to predict EEIs. Overall, the EEI prediction performances of the methods are significantly better than random. The results of our study can further be used by the scientific community in two significant ways.

First, our computational pipelines for predicting EEIs (using the PPDL post-processing) can already be used by the research community. Specifically, the dMaSIF-based EEI prediction pipeline, which shows the best performance with the least computational runtime, can be used on a large scale, as follows. Given a million known protein-protein interaction data (but with no interaction interface information) [51], and a large number of available confident protein 3D structural data because of the recent exponential increase in the human protein 3D structural data of experimental quality [18,19], one can use our dMaSIF-based EEI prediction pipeline to predict EEIs between any two interacting proteins by either using i) one of our pre-trained models or ii) their data to train a model from scratch. Regarding using our pre-trained models, because we use three different EEI definitions, i.e., based on the thermodynamics of the interface formation (D_Engy_), evolutionary conservation of the interface (D_Evol_), or contact-based (D_Con_), and five cross-validation folds, we provide 15 different pre-trained models for our dMaSIF-based EEI prediction pipeline. While the performances of the models vary between cross-validation folds, in general, a pre-trained dMaSIF-based EEI prediction model performs the best for either D_Engy_ or D_Evol_, which should be preferred for predicting EEIs on novel data.

Second, our study provides insights into how to further improve the existing PPIIP methods, designed for the RRI prediction task, in the EEI prediction task. Specifically, when using the RRI prediction scores of the existing PPIIP methods, we show that a convolutional neural network-based post-processing (i.e., PPDL) can learn the patterns in the predicted RRI scores between exon pairs. We show that these learned patterns can then be used to distinguish between interacting vs. non-interacting exon pairs. Looking forward, more sophisticated deep learning-based post-processing approaches might be used to more accurately capture such patterns and thus to further “fine-tune” the existing PPIIP methods for the EEI prediction task.

## Key Points

- We evaluate existing state-of-the-art protein-protein interaction interface prediction (PPIIP) methods, originally designed to predict residue-residue interactions (RRIs), for the exon-exon interaction (EEI) prediction task.
- Using experimentally derived 3D structural data of human proteins, we show that existing PPIIP methods can be extended to predict EEIs, with the best PPIIP method showing AU-ROC values of up to 85% and 70% for the RRI and the EEI prediction tasks, respectively.
- We provide a computational pipeline, incorporating a pre-trained model for the best EEI prediction method, that can be either used to predict novel EEIs in a large scale manner or to train a new model using custom data.

## Authors note

The authors wish it to be known that, in their opinion, Khalique Newaz and Jan Baumbach should be regarded as Joint Corresponding Authors.

## Author Biographies

**Jeanine Liebold** is doctoral researcher at the Institute for Computational Systems Biology and at the Genome Informatics research group at the ZBH - Center for Bioinformatics, Universität Hamburg.

**Aylin Del Moral-Morales** is doctoral researcher at the Universidad Nacional Autónoma de México (UNAM) in the field of Biochemistry. She was a visiting student at the Institute for Computational Systems Biology at Universität Hamburg in 2023.

**Karen Manalastas-Cantos** is a postdoc at the Leibniz-Institut für Virologie, Hamburg. She obtained a joint PhD in Biology from the European Molecular Biology Laboratory and the Universität Heidelberg.

**Olga Tsoy** is a postdoc and group leader at the Institute for Computational Systems Biology at Universität Hamburg. She obtained her PhD from Kharkevich Institute in Moscow.

**Stefan Kurtz** is professor for bioinformatics and head of the Genome Informatics research group at the ZBH - Center for Bioinformatics, Universität Hamburg. He obtained his PhD in Computer Science from Universität Bielefeld.

**Jan Baumbach** is professor and head of the Institute for Computational Systems Biology at Universität Hamburg. He obtained his PhD in Computer Science from Universität Bielefeld.

**Khalique Newaz** is a postdoc at the Institute for Computational Systems Biology at Universität Hamburg.

## Footnotes

1 We use the abbreviation AU-ROC instead of the more common AUC to emphasize that it refers to the receiver-operator curve, in contrast to AU-PRC which refers to the precision-recall curve.

